# Morningness-eveningness assessment from mobile phone communication analysis

**DOI:** 10.1101/2021.02.24.432651

**Authors:** Chandreyee Roy, Daniel Monsivais, Kunal Bhattacharya, Robin I.M. Dunbar, Kimmo Kaski

## Abstract

Human behaviour follows a 24-hour rhythm and is known to be governed by the individual chronotypes. Due to the widespread use of technology in our daily lives, it is possible to record the activities of individuals through their different digital traces. In the present study we utilise a large mobile phone communication dataset containing time stamps of calls and text messages to study the circadian rhythms of anonymous users in a European country. After removing the effect of the synchronization of East-West sun progression with the calling activity, we used two closely related approaches to heuristically compute the chronotypes of the individuals in the dataset, to identify them as morning persons or “larks” and evening persons or “owls”. Using the computed chronotyes we showed how the chronotype is largely dependent on age with younger cohorts being more likely to be owls than older cohorts. Moreover, our analysis showed how on average females have distinctly different chronotypes from males. Younger females are more larkish than males while older females are more owlish. Finally, we also studied the period of low calling activity for each of the users which is considered as a marker of their sleep period during the night. We found that while “extreme larks” tend to sleep more than “extreme owls” on the weekends, we do not observe much variation between them on weekdays. In addition, we have observed that women tend to sleep even less than males on weekdays while there is not much difference between them on the weekends.

## 1 Introduction

Human beings are known to be diurnal in nature that is characterized by a period of activity during the day and a period of inactivity during night. These rhythmic activities are entrained to the light-dark cycles of the solar clock that occurs due to the earth’s rotation around the sun. This rhythmicity is also affected by the social constraints of living in a society, for example going to work on time, while being generally aligned according to the solar clock. The discovery of artificial light has had a considerable impact on the daily activities of human beings, thus eventually affecting their sleeping patterns. However, the physiological activities and behaviour of humans are well known to follow a circadian rhythm that reflects their individual chronotypes.

The chronotype of an individual arises from her natural tendency to align her rhythmic activities according to the solar cycle along with the social constraints of the society. Individuals having early chronotypes rise early in the morning and sleep early as well. They are well known in the literature as “larks”. On the other hand, late chronotypes wake up late as well as sleep late and are befittingly known as “owls”. The rest of the population falls within this spectrum from larks to owls, identified by their individual chronotypes. Identification of chronotypes is an important issue, because an individual’s productivity could depend on the synchrony between her inherent chronotype and her daily work-life timings. We might expect a lark to be more productive during the morning and an owl to be more productive during the evening. Workplaces mostly have schedules that are biased towards early chronotypes and not for the late ones. This can cause sleep deprivation and poor eating habits in the latter which can further lead to health complications [1–4].

Traditionally, studies concerning identification of chronotypes have been done using the Munich ChronoType Questionare (MCTQ) [4–6]. This questionnaire, the first of its kind, consists of unique questions and iconic supporting drawings regarding an individual’s sleep-wake cycle. Survey studies with this type of questionnaire have shown that generally the chronotype of an individual changes gradually with age [7]. Individuals are found to behave like larks in adolescent stages of their lives and gradually tend to become owls through their teenage years reaching a maximum around 20 years of age. After 20, they have been observed to gradually change back to being larks [8]. Additionally, gender differences of chronotypes have also been studied in ref. [9] using the Morningness-Eveningness Questionire (MEQ) [10] in which they concluded the existence of different synchronization patterns for men and women. While these survey studies are excellent tools to understand human behaviour, they have their limitations in the form of sample sizes, memory of the participants, and what is socially and societally expected, etc.

With the advent of the digital age and its rapid development over the years, humans have been increasingly becoming dependent on technology for their daily needs. This has led to them leaving traces of their activity online in the digital world, which in turn can give us considerable insight into their daily activities. Data from mobile phone communication records containing call time stamps and GPS locations along with duration of the voice calls and text messages sent by anonymized users portray periods of activity and inactivity by individuals and consequently are useful for studying their chronotypes. Additionally, data analysis studies harnessing the data from the mobile phone call detail records of a very large number of users presents a very good picture of the dynamics of human behaviour living in a society [11–16]. Access to these kinds of large population datasets enables us to study the social networks formed by humans and relationships formed by them in the networks [17–20]. It can also be used to study migration patterns [21] of the individuals and more recently, it has been used to study the behaviour of people during the COVID-19 pandemic as well [22].

Since mobile phone communication datasets clearly display the circadian rhythms of human activity by looking at the frequency of calls made by an individual during a 24 hour cycle [23], one can broadly determine when an individual is active or inactive. Studies show that the calling activity of individual users on average follow a bimodal distribution where users are active twice during the day with the two peaks in the frequency of calls occurring in the morning and in the evening, respectively. Thus, one can identify the chronotype of an individual by inspecting these rhythmic cycles of calling activity. For example, studies by Aledavood and co-authors [24,25] used data from smartphones of volunteers to identify larks and owls as well as their social networks. They also observed that the personal networks maintained by owls are larger than those maintained by larks.

In our current study, we have used the call detail records of a large population-level dataset to observe the morning and evening calling activities of the users living in a European country during the years 2007, 2008 and 2009. Our aim is to construct a chronotype directly from these activities collected during the entire 24-hour cycle of all seven days of the week instead of only looking at mid-sleep times only on weekends. We observe that they show a bimodal distribution of calling activity that is dependent on the East - West progression of the sun. We have tried to eliminate this effect from the data and have introduced a new approach for the identification of chronotypes of the users by applying factor analysis approaches. First, we carried out a principal component analysis on the data and showed that the first principal component can be interpreted as a chronotype. Furthermore, we used computed values of the chronotype to demarcate between extreme larks, larks, third birds, owls and extreme owls in the population.

Next, we performed an exploratory factor analysis to find if there exist any underlying constructs or factors in our data that govern the behaviour of the individuals. Our study indicates that they behave differently in the morning compared to the evening. We also performed an exploratory bifactor analysis on the dataset which we used to compute a single latent factor that is interpreted to be an individual’s chronotype. This chronotype is a direct manifestation of all the morning and the evening activities of the users on all days of the week and we have shown the variation of the newly computed chronotype with age, along with their gender differences among the individuals. We have observed that the individuals in the younger age cohorts behave as owls whereas older age cohorts tend to behave as larks for both men and women which agrees with previous results found from survey studies [26]. However, men are found to be more owlish than females when they are young and vice-versa when they become older. In addition, through this dataset, one can identify a period of low calling activity, which corresponds to the total sleep duration of an individual in the sleep-wake cycle discussed previously [27]. We have shown that the variation in the duration of sleep is different for weekdays and weekends and is affected by the chronotype for both genders.

## 2 Materials and methods

### Individual mobile phone Call Details Records

The dataset used in this study comprises the Call Details Records (CDRs) from individuals living in a southern European country, which had mobile phone subscription with a specific service provider. It results from the merging of 3 separate subsets (January-December 2007, January-December 2008, and January-December 2009), altogether covering a three year period. The data-sets were anonymised before being handed over by the service provider, such that the true identity of the individual is unknown and each individual is described by a unique identifier (id-number). The CDRs lists all the outgoing calls made by each individual during a three year period, and each entry includes, the id-numbers associated with the caller and the callee, the time and date when the communication event happened, as well as the type of communication event (call or text message) [28]. The data-set includes also user-contract data-sets with some demographic information (age, gender, and registered postal code) of the individuals who were subscribers of the service provider in at least one of the three periods. Over the three year period, different individuals start a new subscription and/or terminate the contract with the service provider, but of the order of six hundred thousand individuals remained loyal to the service provider (*i.e.* their contract started before January 1st 2007 and was still active on December 31st, 2009). From this set of loyal subscribers whose demographic information was available (some user have missing entries or they contain typos), we chose 11178 individuals, who made at least 100 hundred calls/text messages each year and with a total number of calls/sms not exceeding 5000 calls (to exclude possible calls centers or subscription sellers).

### Geographical grouping based on individual’s location

Using the information of the postal code available in the user-contract files, we split individuals into 5 groups, each one falling inside a longitudinal band enclosing their geographical location. However, as a signed non-disclosure agreement (NDA) prohibit us to disclose the country where the service provider offered, the actual values delimiting each selected geographical longitudinal band are masked. From here on, the longitudinal values will be reported from a reference point located near the easternmost part of the country, which will work as the zero reference. Thus, five latitudinal bands *L_I_*-*L_V_* of widths 2.5°, 3.05°, 2.75°, 2.2° and 2.8° are defined, separated by exclusion bands of width 0.2 degrees, and with the first band *L_I_* (the reference point) the easternmost, and the subsequent bands located to the west progressively until the last longitudinal band *L_V_* lies in the westernmost part of the region (13.3°-wide). Table 1 lists the number of individuals in each longitudinal band, as well, gender and age distribution information of the population on each band.

**Table 1.** Demographic information of the population

| Feature | Longitudinal Band |  |  |  |  |
| --- | --- | --- | --- | --- | --- |
| | $L_I$ | $L_{II}$ | $L_{III}$ | $L_{IV}$ | $L_V$ |
| Limits | $[0^\circ, -2.3^\circ]$ | $[-2.5^\circ, -5.35^\circ]$ | $[-5.55^\circ, -8.1^\circ]$ | $[-8.3^\circ, -10.3^\circ]$ | $[-10.5^\circ, -13.1^\circ]$ |
| Width | $2.5^\circ$ | $3.05^\circ$ | $2.75^\circ$ | $2.2^\circ$ | $2.8^\circ$ |
| Individuals | 2031 | 2589 | 2386 | 2816 | 1356 |
| Females | 1108 | 1479 | 1338 | 1564 | 795 |
| Males | 923 | 1110 | 1048 | 1252 | 561 |
| Young (18-35) | 637 | 891 | 854 | 928 | 443 |
| Mid-age (35-60) | 1017 | 1272 | 1177 | 1425 | 700 |
| Old (60-78) | 377 | 426 | 355 | 463 | 213 |

The reason for this longitudinal splitting is to take into account of the dependence of the human chronotype on the East-West progression of the Sun, as has been shown in earlier studies [27,29]. In Fig 1 plots, for each of the five longitudinal bands *L_I_-L_V_*, the calling activity of the analyzed individuals during weekday nights (Mondays to Thursdays) aggregated over the 3 year period. There, a clear shift between the calling activity distribution of each region can be seen, with the easternmost band *L_I_* starting and ending its calling activity around 45 minutes earlier than the westernmost band *L_V_*.

**Fig 1.**
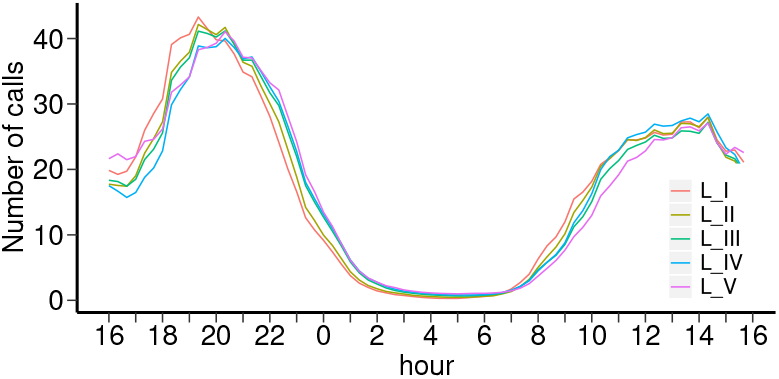
Aggregated individual calling activity for weekdays in 5 geographical regions. For each the 5 longitudinal bands *L_I_-L_V_*, the calling activity of the analyzed individuals for all 24-hour over-the-night periods during weekdays (Mondays to Thursdays) aggregated over the 3 year period. The activity in the easternmost band *L_I_* (red line) is noticeably shifted towards early hours compared to the activity in the westernmost band *L_V_* (magenta line). The bands limits are as follows: *L_I_*: [0°, −2.3°], *L_II_*: [−2.5°, −5.35°], *L_III_*: [−5.55°, −8.1°], *L_IV_*: [−8.3°, 10.3°], and *L_V_*,: [−10.5°, −13.1°].

## 3 Mid-sleep time and sleep duration

In order to determine individual’s daily periods of non-activity, we analyze separately the number of events (calls/text message) that each individual made on different days of the week for over the 3-year period. As we are interested in determining the mid-sleep time, we determine the calling activity taking place on each night of the week, such that we split it in seven 24-hour periods each one starting at 4:00pm (*e.g.* 4:00 pm Monday, and ending at 3:59pm on the next day *i.e.* Tuesday). From here on, we refer to these periods as “nights”, with, for example, Saturday’s night meaning the time period between Saturday 4:00pm and Sunday 3:59pm. In addition, the first four nights of the week (Monday to Thursdays) are also aggregated into a 24-hour long period named “weekday night”, which is a standard way to refer to in chronotype studies to workdays.

Using these definitions, we aggregated the weekly calling activity of each individual over the 3-year period on the corresponding period of the week (Weekday, Friday, Saturday and Sunday nights). In Fig. 2, we show the aggregated calling activity of one individual for the four night periods studied.

**Fig 2.**
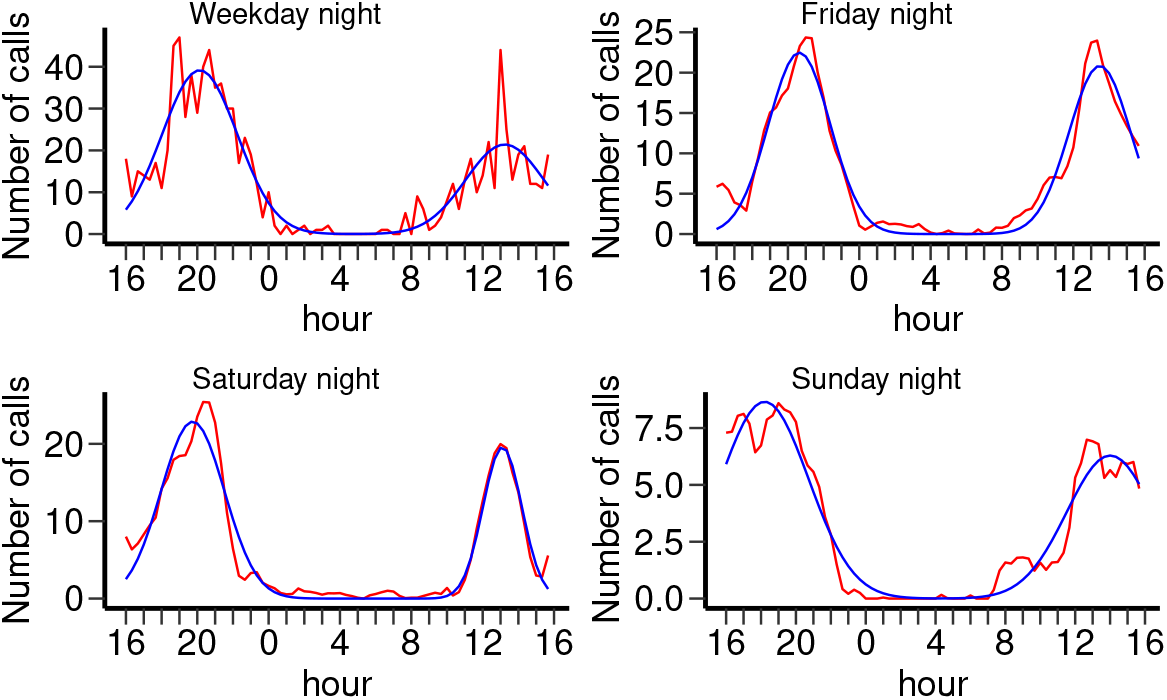
Aggregated calling activity of one individual for 4 different overnight periods. The calling activity as a function of the time of the day (red lines), for four different overnight periods: Weekdays (top-left), Fridays (top-right), Saturdays (bottom left) and Sundays (bottom-right). Blue lines are smoothed Gaussian curves. For this individual, the minimum of activity typically occurs around 5:00 am.

The bimodal distribution shown in Fig. 2 for one individual is present in almost all the individuals’ profiles, and the consistent bimodality shown in the average calling activity of the population at 5 different longitudinal bands (see Fig. 1) is a reflection of this generality. In Fig. 3 we plot the calling activity patterns of a sample of users in one of the 5 latitudinal regions (arranged in an actigram-like representation) to show that the bimodality of the daily overnight calling activity is consistent. This bimodal pattern will be used to approximate each individual’s calling activity by a Gaussian Mixture Model (GMM) [30], which has recently been used to describe human activity from CDRs [23,31].

**Fig 3.**
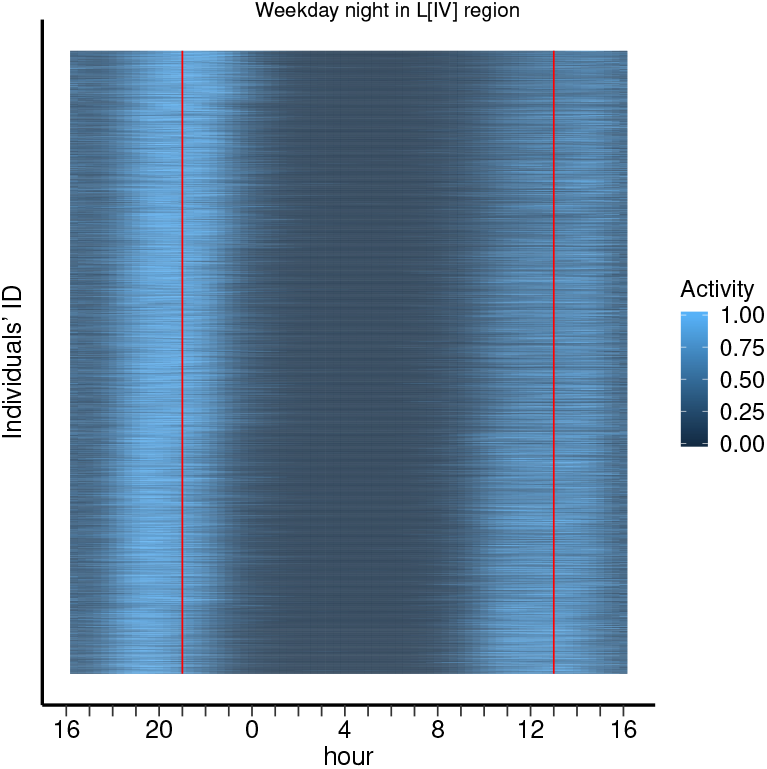
Actigram of individual calling activity during weekdays. An example of an actigram showing the calling activity for weekdays’ overnight periods of 1560 individuals chosen from the longitudinal band *L_IV_* (similar patterns exist for the other bands). Each individual calling activity correspond to a horizontal line in the actigram, and each individual activity is scaled into the interval [0,1] such that the most active periods are represented by light regions and periods without activity by dark regions. To show that individuals’ period of low activity (chronotypes) are not homogeneous, we ordered the presented individual activity profiles according to time shift between each profile and the mean over the population’s activity profiles (*i.e.*, using the time shift that maximizes the cross-correlation between the individual profile and the mean). In this representation, “larks” or morning people) appeared in the bottom section of the actigram, with their activity profile clearly shifted towards early hours, whereas “owls” or evening people (top section of the plot) are clearly shift towards late hours.

A GMM with two modes (Gaussians) used as an approximation of the calling activity is given by:

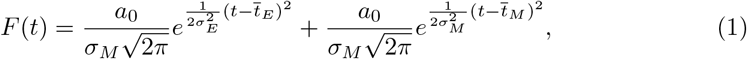

where 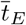 and *σ_E_* are the mean and the standard deviation of the Gaussian located in the evening (left) and 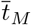 and *σ_M_* the corresponding values for the one located in the morning (right) of the following day.

The means 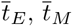 and the standard deviations *σ_E_, σ_M_* given by the approximations can be used to describe the relevant quantities of each individual activity pattern, namely the sleeping duration *T_LCA_* and the mid-sleep time *T_mid_*. Assuming that the period of sleeping is bounded by the period when the calling activity falls to a minimum, we can approximate the sleeping duration or the period of low calling activity *T_LCA_* of the day of the week *d* by the width of the area between the activity modes, that is,

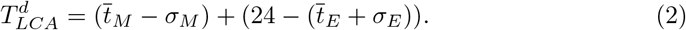

Similarly, the mid-sleep time 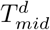 of the day of the week *d* is taken as the midpoint between the calling activity modes, thus

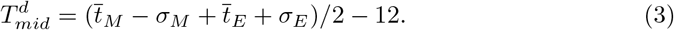

### 3.1 Morningness-Eveningness classification

The means 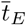 and 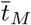, and the standard deviations *σ_E_* and *σ_M_* calculated using Eqn. 1 are not always well defined because of the randomness of individuals’ calling activities. Therefore, after filtering out the outliers in the dataset we consider a total of 11178 individuals for our analysis with 2031, 2589, 2386, 2816 and 1356 in longitudinal bands *L_I_, L_II_, L_III_, L_IV_* and *L_V_*, respectively. The definitions of the weekly overnight periods (Weekday-, Friday-, Saturday-, and Sunday-night) have associated same number of mid-sleep times 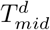, which can be determined for each individual from the calling activity. These four mid-sleep times can be used to assess the tendency of an individual to have early (morningness person) or late (eveningness person) schedules.

In general, the mid-sleep times of any individual depends on the day of the week, such that the mid-sleep times on weekdays occur usually earlier than on weekends. However, when comparing between individuals, one can expect that the set of mid-sleep times from a morning person, occurs in general earlier than those of an evening person, thus we can use this expected difference for chronotype classification. The correlations between 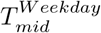 and 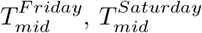 and 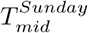 are 0.65, 0.54 and 0.67, respectively, which corroborates that in general individuals having early schedules on weekdays have also early schedules on weekends.

In spite of the differences found between the different chronotypes that each individual has for different days of the week, the individual has a consistent type (morningness or eveningness) relative to other individuals in the population. Comparing the chronotypes between individuals, those having earlier schedules on weekdays have also earlier schedules on weekends, and similarly for those having later schedules. We use this consistent order between daily chronotypes between individuals to assess their morningness-eveningness. The four possible mid-sleep times (on Weekdays, Fridays, Saturdays and Sundays)are assigned to a 4-dimensional vector

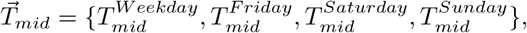

and this vector will be used to assess the chronotype.

Next we apply Principle Component Analysis (PCA) in the space of vectors 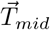 of the population to get a better representation of the chronotype vectors. The loadings of the 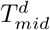on the first principal component (**PC1**) has been provided in the SI. We have plotted a summary of the negative of the **PC1** (see S1 Table) in a box-plot on the left in Fig. 4A for the populations in the five longitudinal bands to exhibit the East-West progression of the mean values. A positive **PC1** can be interpreted as an individual having a later mid-sleep time and a negative value can be interpreted for her to have an earlier mid-sleep time. These values can thus be used to understand the morningness-eveningness of an individual in the population. We observe in Fig. 4A, that the mean of the **PC1** decreases from *L_V_* to *L_I_*, which would imply that there are more larks in the Eastern part of the country than in the Western part, which, however, could be misleading [27]. In order to remove this possible artefact, we have computed a multiple linear regression model of 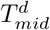 values with latitude and longitude of the users as independent variables. The coefficients of the latitude and longitude computed from the model along with their p values has been summarized in S2 Table. We observe that the longitude is most significant for all 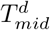 while there is very small dependence on latitude. Therefore, we considered the residuals computed from the regression model and have again applied a PCA on them. On the right of Fig. 4A, we have shown a summary of the new first principal component (**PC1**_*reg*_) for the vector 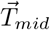 in a box-plot which shows that the effect of East-West progression has been removed.

**Fig 4.**
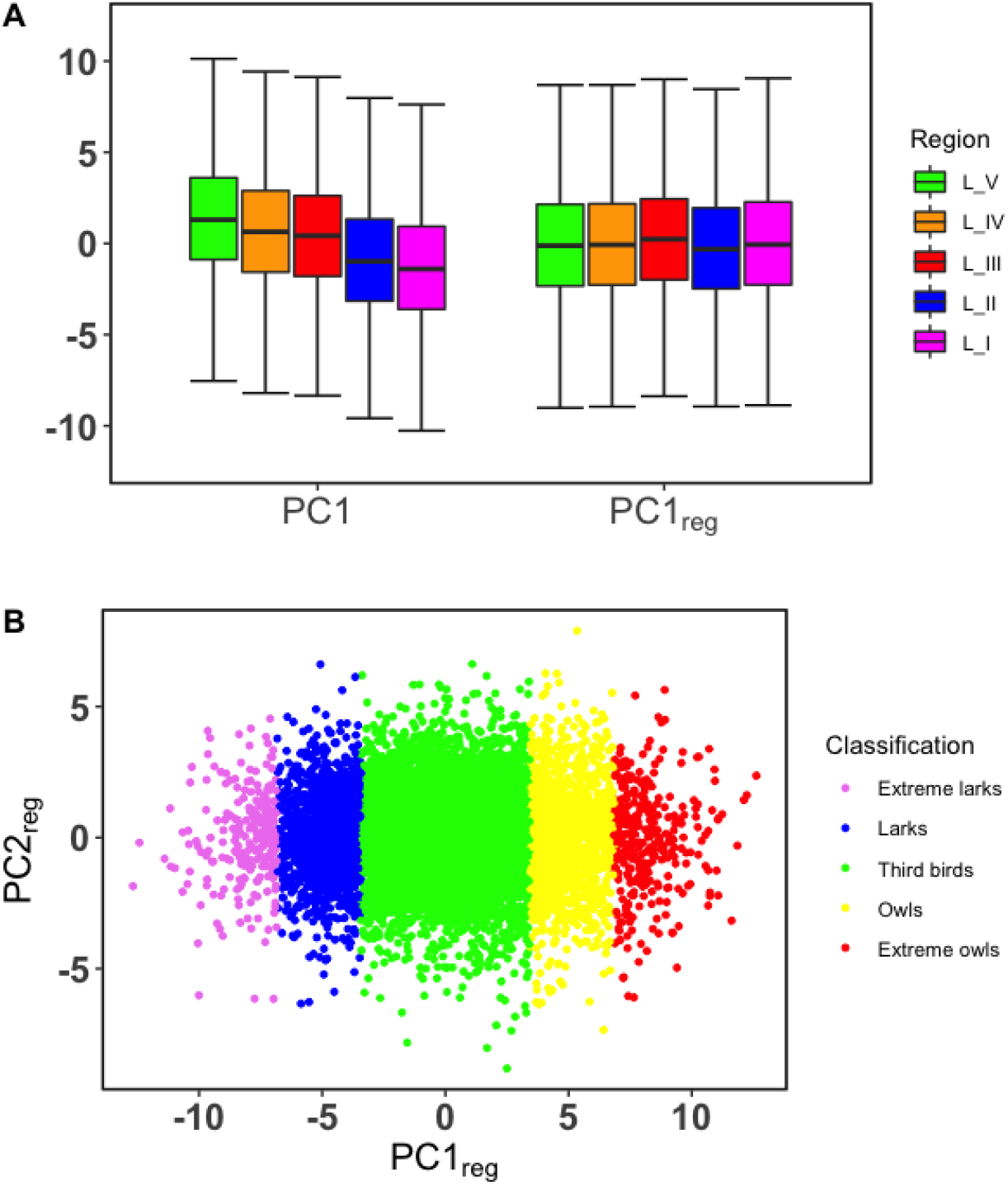
Principal component analysis (PCA) on the four mid-sleep times 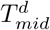. (A) On the left a summary of the first principal component (**PC1**) obtained from PCA computed on the four sets of 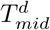 for all users in five different longitudinal bands of the country. On the right, we have shown a summary of the first principal component (**PC1**_*reg*_ or p chronotype) obtained after applying PCA to the residuals obtained from a regression analysis of 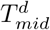 using the latitude and longitude as independent variables. The green, orange, red, blue and magenta bars represent the westernmost, western, middle, eastern and the easternmost parts of the country, respectively. The black horizontal lines in the middle represent the median of the distribution. The box plot includes all the values within the range of the 25^*th*^ and 75^*th*^ percentile and the end of the whiskers represent the maximum and the minimum scores excluding outliers. (B) The first two principal components (**PC1**_*reg*_ and **PC2**_*reg*_) of the PCA on regressed values of the chronotypes in vector 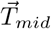. The distribution of the **PC1**_*reg*_ or p chronotype is slightly skewed (skewness = 0.12), and leptokurtic (kurtosis = 3.09). Therefore, one can divide the individuals into five groups using the mean and standard deviation of the distribution called extreme larks, larks, third birds, owls and extreme owls according to their chronotype and are represented in the figure by the colours violet, blue, green, yellow and red, from left to right, respectively.

Once the set chronotype vector (**PC1**_*reg*_ or p chronotype) has been transformed under the PCA, we observe that the resulting distribution of its values has a small skewness of 0.12 and is very slightly leptokurtic with a kurtosis value of 3.09, in comparison with the Gaussian distribution having skewness 0 and kurtosis 3. Since the distribution is positively skewed, we expect there to be slightly more owls than larks present in the population. The individuals can be divided into five clusters, inline with the standard classification in the literature of the morningness-eveningness into five groups (definitely morning-type, moderately morning, neither-type, moderately evening-type, and definitely evening-type) [9]. In Fig. 4B we have divided the users into these clusters using the means (*m*) and standard deviation (*σ*) of the distribution of **PC1**_*reg*_. Hence, the individuals divided by the values: *m* – 3*σ, m* – 2*σ, m* – *σ,m* + *σ,m* + 2*σ, m* + 3*σ*, are accordingly named as extreme larks (violet), larks (blue), third birds (green), owls (yellow) and extreme owls (red), and comprise 2.22%, 13.05%, 69.04%, 12.84%, and 2.85% of the population, respectively.

Furthermore, we have computed a PCA on the *T_mid_* values for the weekdays and weekends separately to observe changes in the behavioural traits of larks and owls. The first principal component obtained from PCA on mid-sleep times on weekdays 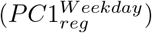 accounts for 83.7% of the variance in the data and the one obtained from PCA on mid-sleep times on weekends 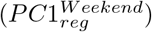 account for 81.8% of the variance in the data. The distribution of the PC1s obtained are more leptokurtic and more skewed than the distribution of the p chronotype. 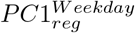 and 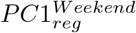 are observed to have a kurtosis of {3.26, 3.25} and skewness of {0.18, 0.16}, respectively. One can then classify the five different groups of people (extreme larks, larks, third birds, owls, extreme owls) using the same method described in previous paragraph. In Fig. 5 we show a Venn diagram that depicts the joint distribution of the PC1 distributions considering only the larks and the owls. Here we observe that approximately 7. 1% of larks and 7.4% of owls of the total population show the same behavioural traits on both weekdays and weekends. Around and 8% and 7.4% of larks on weekdays and weekends respectively change to third birds. Similarly 7.6% and 8% of owls on weekdays and weekends, respectively, convert to third birds. A very small percentage of the population, 0.1% of the larks and 0.3% of the owls, on weekdays and weekends change to opposite behaviour, i.e. larkish become owlish or owlish become larkish.

**Fig 5.**
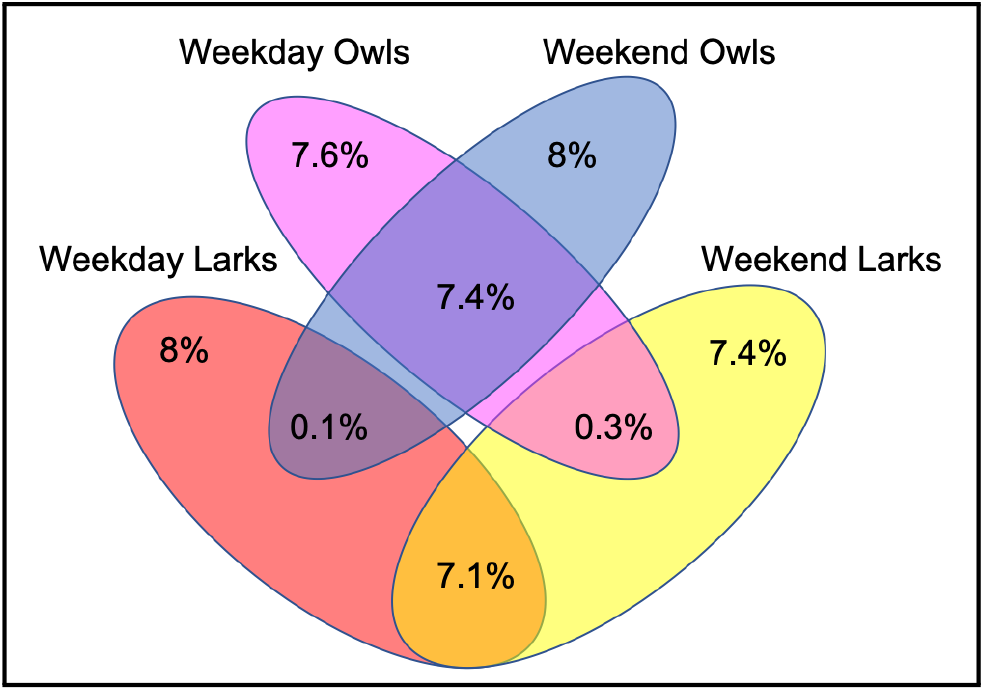
Venn diagram to show changes in larkish and owlish behaviour. The joint distribution of the larks and owls obtained from classification of the chronotypes obtained by computing a PCA on *T_mid_* values for weekends and for weekdays separately and presented in the form of a Venn diagram. We observe that 7.1% of the population who are larks remain the same be it weekdays or weekends and similarly for 7.4% of the population that are owls. A very few percentage of the population (0.1% for larks and 0.3% for owls) changes their behaviour drastically to the opposite kind depending on whether it is a weekend or weekday. We also observe that 8% for larks and 7. 6% for owls on weekdays; and 7.4% for larks and 8% for owls on weekends convert to third birds in the population.

## 4 Model for morningness-eveningness assessment using factor analysis

In general, the individual mid-sleep times are different on different days of the week, with the earliest mid-sleep time occurring in weekdays and the latest on Saturdays (around one hour difference on average). When comparing mid-sleep times between individuals, there we observe the following. The relative order between individuals is, in general, the same regardless of the period of the week analysed, such that individuals belonging to the group with earlier chronotypes on weekdays (relative to the whole population), also belong to the groups with earlier chronotypes on weekends.

In this section, we have attempted to compute a chronotype score using factor analysis for all the users in the population that can reflect the morningness or eveningness of an individual. The first maximum in the activity shown in Fig. 1, considering the night centered approach, represents an average peak in the evening activity (EA) of a user on a particular day of the week and the second peak represents the morning activity (MA). These pairs of observables for each day have been computed from the data for each individual and is denoted by MA*d* and EA*d* where d stands for Weekdays, Fridays, Saturdays and Sundays, respectively. Generally, the MAs of individuals are mostly constrained due to social obligations, like going to work on time. In the evening, they are more relaxed and can follow their own individual chronotypes. Therefore, we hypothesize that the morning and evening behaviour of an individual are different from each other and these can be used to obtain their chronotype. We have considered an Exploratory Factor Analysis (EFA) [32, 33] on the sets of observables for {MA*d*} and {EA*d*} as shown in Fig. 6A to explore an underlying latent variable that affects an individual’s morning and evening activities.

**Fig 6.**
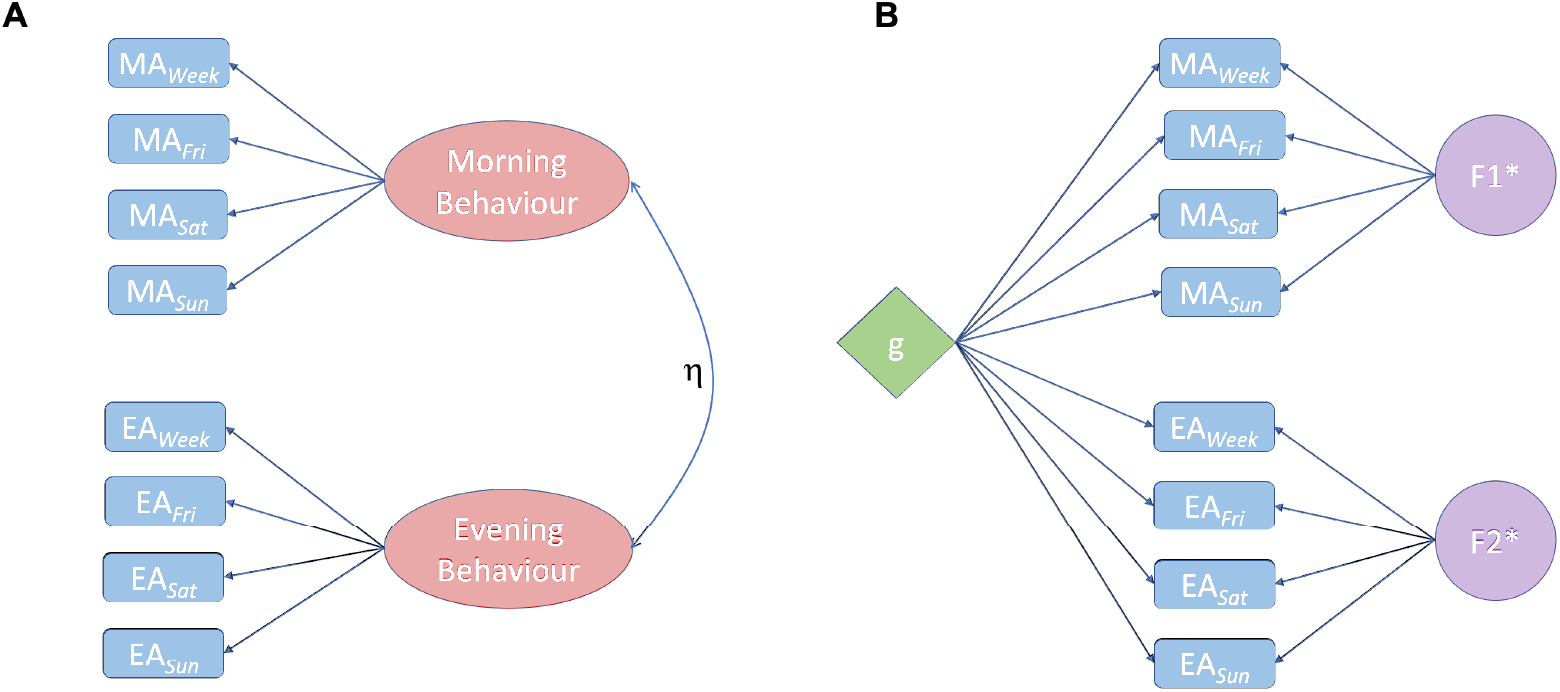
Factor analysis of the chronotype of an individual using CDRs: (A) An Exploratory Factor Analysis (EFA) of the peak locations of the morning activity (MA) and evening activity (EA) on the seven days of the week shows the emergence of two underlying latent factors (Morning behaviour and Evening behaviour of an individual) that governs the outcome of MA and EA, separately. The correlation between the two latent factor is computed to be *η* = 0.31. (B) A higher order model (Bifactor model) is used to find an underlying general g factor that can be used to understand the chronotype of an individual. *F*1* and *F*2* are the “group” factors that affect the MAs and EAs, separately. The loadings of the factors on the observables for both the figures have been specified in two separate tables (S3 Table and S4 Table) in the SI. Any cross-loadings with values below 0.3 have been ignored.

We first assess the factorability of the data by carrying out the following tests. The Kaiser-Meyer-Olkin test for factor adequacy in the model gives a score of 0.74 [34]. In addition we use the Bartlett’s test for sphericity [35] to check any redundancy between the observables that are summarized with fewer number of the latent variables and it was gives a χ^2^ = 32344.32 and *p* < 0.0001. Both tests indicate a favourable use of EFA in the model. The EFA using oblimin rotation was conducted using “psych” package’s [36] maximum likelihood method to extract the factors. All items have high communality value around 1.0 indicating that the two factor structure in the model is able to explain a large part of the variances in the observables. This agrees with our aforementioned hypothesis and accordingly we have observed that all the MAs are loaded on one factor (Morning behaviour) and all the EAs are loaded on another factor (Evening Behaviour) as depicted in Fig. 6A. The correlation between the two factors is *η* = 0.31. This model has a unidimensionality score of 0.59 which further supports our claim that the individuals behave differently in the morning from that in the evening. The individual factor loadings on the observables are summarised in S3 Table in the SI. The cross-loadings of the factors have not been shown in this figure since their values are less than 0.3. The EFA is a technique used to identify conceivable underlying constructs within the observables and is distinct from PCA that is used to reduce the dimensions in the data. The model fit indices for the EFA, namely, comparative fit index (CFI), Tucker-Lewis Index (TLI), and the root mean square error of approximation (RMSEA), have been computed to be 0.91, 0.80, and 0.14, respectively. While the values of CFI and TLI indicate a good fit of the data in our model, we get a high value for the RMSEA.

Next, we have carried out an exploratory bifactor analysis (EBA) [38] on the model to determine scores for a single construct like the chronotype of an individual that would reflect the morningness or eveningness of the person even when the data is multidimensional. The bifactor models are useful in representing hierarchical latent structures in the data as the first-order factors [39]. It computes the factor scores for a general factor *g*, which loads directly onto all the observables in the model and also produces group factors that distinguish between the groups formed among the observables. We have used “omega” function from the package “psych”, which does a factor analysis followed by an oblique rotation and extracts the general factor using Schmid-Leiman transformation [40,41]. The tests of reliability in our model *ω_t_* and *ω_h_* are computed to be 0.84 and 0.39, respectively. Here *ω_t_* accounts for the total variance in the data due to the general factor g and the group factors together, whereas *ω_h_* accounts for the proportion of variance in the data due to the general factor only. In Fig. 6B we show that the all the observables or items are loaded on the general factor (*g*), which represents the chronotype of an individual. The factors represented by F1* and F2* are group factors or *nuisance dimensions—factors* that measure responses of the observables that are not considered by the *g* factor. The loadings of all the factors in this analysis have been summarized in S4 Table in SI.

The mean factor scores obtained for the morningness behaviour, eveningness behaviour, and the g chronotype from the models discussed above are found to behave in a way similar to the principal component in the left of Fig. 4A when plotted as a function of the longitudinal bands from West to East. As discussed previously in the case of PCA we have computed a multiple linear regression model of the observables with the latitude and longitude of the users as independent variables (details in S5 Table of SI) to remove the geographical dependence in the data. We have considered the residuals computed from the regression model for all the items in our analysis. We have applied EFA and EBA on the residuals and the corresponding plots are shown in S6 Fig in SI.

### 4.1 Age and gender dependence of the chronotype

The factor scores obtained from our models can be interpreted as an indicator of a user’s chronotype. Individuals having negative scores are considered to be larkish or morning-type and those having positive scores to be owlish or evening-type. The higher the value of the scores, the more extreme larkish or extreme owlish behaviour of an individual is expected to be. Fig. 7 shows the average factor scores of (A) Morning Behaviour - MB, (B) Evening Behaviour - EB and (C) the g chronotype as a function of the users’ age and gender. The factor scores for g chronotype in Fig. 7C for both the genders are found to be decreasing with age indicating that younger individuals (from 18 to 35 years old) are more owlish in nature. Furthermore, it is also observed that males in the younger age cohorts tend to have higher factor scores than women indicating that they are more owlish than females. However, after 35 there is a crossover and the females are observed to be more owlish in nature than males. For the mid age cohorts (between 35 to 60 years old), we observe a peak in the factor scores. Finally, older age cohorts (above 60 years) are found to behave like larks with women still having later chronotypes than men. The age and gender dependence of the p chronotype computed using PCA discussed in a previous section has also been shown in Fig. 7D. The variation of p chronotype is observed to be qualitatively similar to the one observed with g chronotype. We have also observed a high correlation between the two chronotypes and it is found to be 0.76.

**Fig 7.**
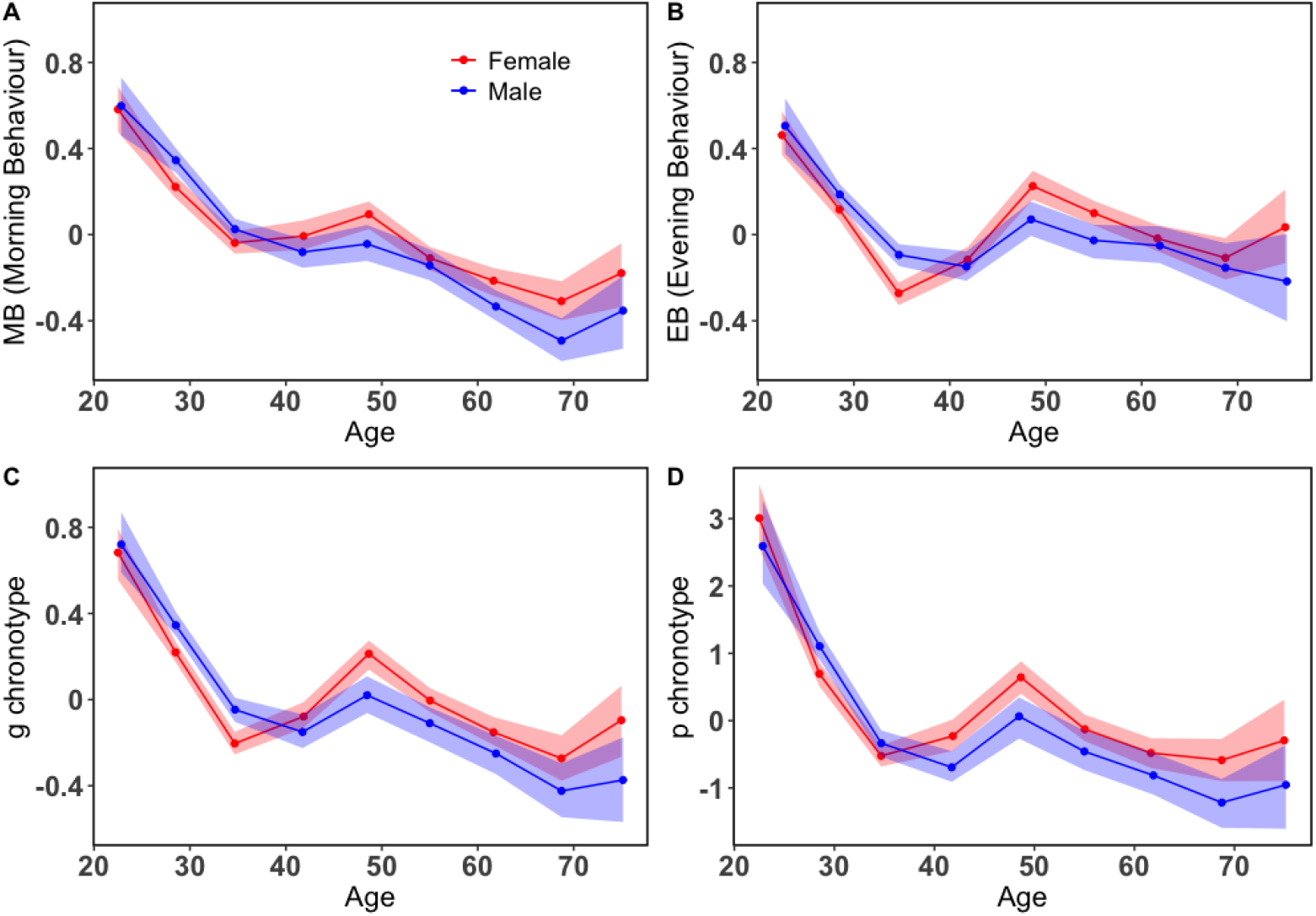
The mean of the factor scores and first principal component computed from PCA for different age and gender cohorts: The variation of the average factor scores of the (A) morning behaviour, (B) evening behaviour and (C) the general g chronotype along with (D) the first principal component (**PC1**_*reg*_) from the PCA renamed as p chronotype is plotted as a function of age and gender. Each point is calculated separately with the shaded regions in all the three plots representing the bootstrap 95% confidence intervals. The curve in red is for females and blue for males.

### 4.2 Dependence of the period of low calling activity on the chronotype

The period of low calling activity *T_LCA_* which can be interpreted as a representation of an individual’s sleep duration during the night time, has been calculated using Eqn. 2. Furthermore, we have calculated the average sleep duration on weekend (*T_LCA_* – *Weekend*) and weekdays (*T_LCA_* − Weekday) separately and Fig. 8A shows their variation as a function of the g chronotype. We find that the users sleep more on the weekends than on weekdays and this result is consistent with the previous findings [42]. Moreover, larks are observed to sleep more than owls on weekends. This is because both larks and owls tend to align themselves according to their own chronotype as there are no social constraints governing their schedules. Since the larks tend to follow the solar clock they tend to sleep more than owls. On weekdays, extreme owls have the same sleep duration as the larks which implies that they are not able to keep up with the social constraints like work schedules and end up oversleeping. Fig. 8B shows the sleep duration on weekends as function of the chronotype and the gender. We do not observe any significant differences between the two genders on the weekends. However, in Fig. 8C we find that owlish males sleep more on weekdays than owlish females. The average age of the females in this regime falls in the age cohort of 40 to 60 year old. This could be a reason for the less sleep duration as they are more active than males around this age due to reasons already discussed in the previous section.

**Fig 8.**
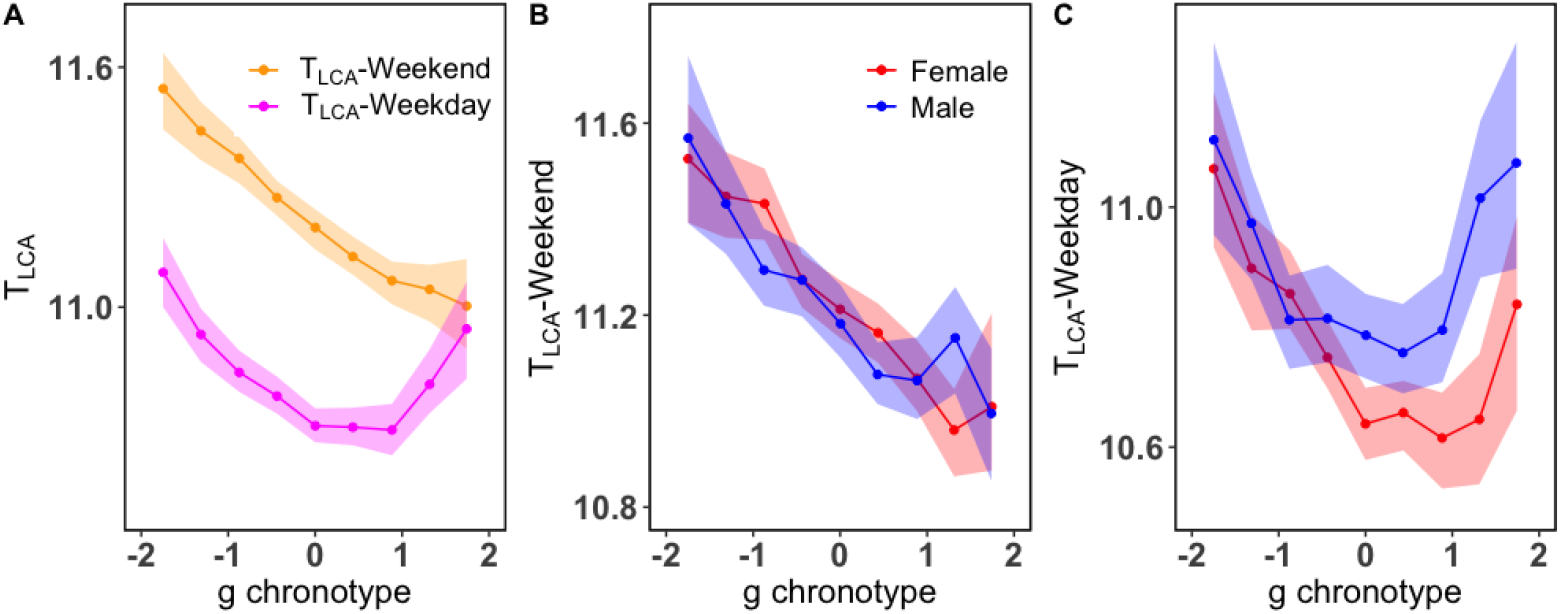
Period of low calling activity of users of different chronotypes: (A)The period of low calling activity 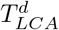 is the width of the area between the activity modes, calculated according to Eqn. 2 for *d* different days of the week. The average of 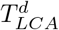 on weekends *T_LCA_* – Weekend and weekdays *T_LCA_* – Weekday is plotted as a function of the general *g* chronotype in orange and magenta, respectively. (B) *T_LCA_*-Weekend and (C) *T_LCA_* – Weekday are plotted as a function of the chronotype for both genders. For each chronotype, *T_LCA_* Weekend and *T_LCA_* Weekday has been calculated separately with the shaded regions in all three plots representing the bootstrap 95% confidence intervals. The red plot is for females and blue for males in both (B) and (C).

## 5 Discussion

In this study, we have utilized mobile phone communication data of a population in a European country to study chronotypes of the service users. We have shown that the chronotype of an individual can be broadly identified through two related yet distinct statistical approaches. In our first approach, we have used PCA to heuristically calculate a composite score for the different chronotypes using the mid-sleep times on weekdays along with the weekends as well. The first principal component from this analysis is composed of all the mid-sleep times with all positive loadings. We observed a slightly leptokurtic distribution with a small skewness for the computed values. Using the mean and the standard deviation of the distribution, we divided the users into the following five different clusters - third birds, the users in the centre (69.0%), larks (13.1%), owls (12.8%), extreme larks (2.2%), and extreme owls (2.9%). Moreover, if one were to do the PCA for weekdays and weekends separately, the joint distribution obtained from the two first principal components indicate changes in the behavioural traits of the individuals. While some of the larks and owls behave the same on weekends and weekdays, some have also been observed to change to the third birds category. We also observed very small percentages of larks to convert to owls and vice versa.

Using a second approach, i.e. the EFA, which assumed chronotype to be a latent trait, we also found that the morning activities and evening activities of the users are governed by two separate factors, namely the morning behaviour and the evening behaviour. The morning behaviour of the users is usually more constrained due to the society following a strict schedule for offices, schools, etc. and most chronotypes try to align themselves accordingly. However, this is not the case in their evening behaviour since the individuals are more flexible with their evening schedules. Therefore, we see a more pronounced change in the morning behaviour than in the evenings as depicted in Fig. 7A and B. Furthermore, it is seen that older cohorts usually do not follow a strict schedule since they are mostly retired from work and consequently tend to follow their inherent chronotypes. In contrast, the younger cohorts need to follow a stricter social timetable and thus exhibit a vastly different behaviour than the older cohorts.

Traditionally, the chronotypes have been calculated using the mid-sleep time of the individuals on weekends only since they are assumed to follow their inherent chronotypes freely during these days of the week [43]. However, in our study we have shown that the g chronotype computed using an EBA is also an appropriate method to study the morningness and eveningness of an individual. It is a higher order version of the EFA that is able to compute a general factor g that is directly related to all the observables in the data. Hence, this general factor renamed as g chronotype from EBA is able to capture all the effects of the activities of an individual on all days of the week. We found that the younger cohorts tend to have later chronotypes, which gradually change to earlier chronotype with increasing age, similar to the results of the earlier survey study [43]. However, the variation in the chronotype reduces considerably for age groups above 40. In addition, we have observed a small peak for mid-age cohorts that could be a direct influence of the lifestyle led by most of the individuals in this age group. Most of them have to connect with both their children (young age cohorts) and parents (old age cohorts) who usually live in separate accommodations. Thus, the peak can be assumed to be a direct manifestation of their calling activity needed to maintain their social interactions with both the age groups. The individuals in the older age cohorts (60 years and above) are most likely not adhering to regular work schedules and so, they tend to follow their inherent chronotype, which is more aligned with their biological and the solar clock. This could be a reason for them to follow a more larkish behaviour. These changes can also be attributed to other factors like hormonal changes during an individual’s life span that affects their sleeping patterns [8]. Additionally, women above the age of 40 are found to show more owlish chronotype when compared to men. This trait may be a direct cause of societal responsibilities that are usually assumed to be taken predominantly by women e.g. child care etc [44].

Furthermore, we have observed similar behaviour for both the g chronotype and the p chronotype computed from EBA and PCA, respectively, as depicted in Fig 7D. Chronotype m computed from an EFA on mid-sleep times for all days is found to show the same variation as g chronotype (see S7 Fig in SI). Using the g chronotype we are able to demonstrate that the chronotypes identified by directly taking into account and combining several observables of human activity, instead of a derived quantity like the mid-sleep time, can also be used to distinguish between the morningness and the eveningness of individuals. Moreover, our results agree with the previous findings using traditional methods like the MCTQ and MEQ questionnaires [8,29,45–48]. Using a period of the users’ low calling activity as markers of their sleep duration [27] we find that on average all chronotypes sleep more on weekends than on weekdays [42] and in both cases larks are generally found to sleep more than owls. On weekends, larks go to sleep earlier than owls and so they have a longer sleeping period. The shorter sleeping periods observed for owls may be a cause for sleep deprivation occurring among them, which can further lead to health issues [1–3]. However, on the weekdays we observe that extreme owls have sleep duration similar to extreme larks suggesting that on weekdays the former may have difficulties in observing work schedules.

Finally, we conclude that our results obtained by combining data from mobile phone communication of individuals during a 24 hour day-night cycle, one can form a detailed understanding of their chronotypes. These kinds of studies using mobile phone service subcribers’ CDRs, demographic, and location information gives a novel and time-wise longitudinal perspective to the circadian rhythms of individuals. Our data-driven approach adds and complements the questionnaire based studies and findings in them, as it avoids possible shortcomings in terms of sample size and dependence on the memory of the participants. In the recent past, the rapid adoption of newer modes digital communication and different smart devices have largely supplemented the usage of mobile phones. Therefore, the approach of combining digital data from multiple channels of communication to assess the chronotype of an individual as a reflective or a latent trait would be extremely worthwhile and timely in fields such as mobile health and medicine [49, 50].

## Acknowledgments

CR, DM, KB, and KK acknowledge support from EU HORIZON 2020 INFRAIA-1-2014-2015 program project (SoBigData) No. 654024 and INFRAIA-2019-1 (SoBigData++) No. 871042. KK also acknowledges the Visiting Fellowship at The Alan Turing Institute, UK.

## Supporting information

**S1 Table.** Principal component analysis of the mid-sleep times. We have performed a principal component analysis on the mid-sleep times of the vector 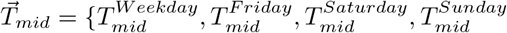 as discussed in the main text. The loadings of the 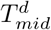 has been summarized in a table. Since all the loadings have negative values we use -**PC1** as a convention to study the chronotypes from 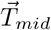. The reversal of sign does not affect the results in the case of PCA because the principal axis is rotated arbitrarily to get the best fit of the data. Moreover, we have found that the correlation between the PC1 and g chronotype (discussed in details in later sections) have a negative value of −0.73. Thus we have reversed the sign for non regressed values of 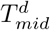 to maintain the same conventions for all chronotypes.

| | $T_{mid}^{Weekday}$ | $T_{mid}^{Friday}$ | $T_{mid}^{Saturday}$ | $T_{mid}^{Sunday}$ |
| --- | --- | --- | --- | --- |
| PC1 | -0.47 | -0.50 | -0.54 | -0.50 |

**S2 Table.** Regression of the mid-sleep times. To remove the effect of the East-West sun progression, we performed a regression on the 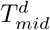 values for all d days with latitude and longitude as independent variables. The values of the intercept, coefficients of latitude and longitude have been enumerated in this table along with their p values.

|  | Intercept | p value | Longitude | p value | Latitude | p value |
| --- | --- | --- | --- | --- | --- | --- |
| $T_{mid}^{Weekday}$ | 35.45 | < 0.0001 | -0.12 | < 0.0001 | 0.02 | 0.005 |
| $T_{mid}^{Friday}$ | 38.14 | < 0.0001 | -0.14 | < 0.0001 | -0.01 | 0.114 |
| $T_{mid}^{Saturday}$ | 38.43 | < 0.0001 | -0.16 | < 0.0001 | 0.0 | 0.786 |
| $T_{mid}^{Sunday}$ | 35.64 | < 0.0001 | -0.11 | < 0.0001 | 0.04 | 0.0004 |

**S3 Table.** Exploratory factor analysis. An exploratory factor analysis performed on the morning and evening activities of the dataset reveals that there are two underlying constructs or latent factors in the data. They are characterized by morning behaviour and evening behaviour of the individuals and the loadings of these factors on the observables i.e. the calling activities of the individuals have been summarised in this table. The boxes coloured in blue shows the high values of the MB on morning activities only and EB on evening activities clearly illustrating the two distinct factors arising from factor analysis.

|  | Morning Behaviour (MB) | Evening Behaviour (EB) |
| --- | --- | --- |
| $MA_{Work}$ | 0.90 | -0.02 |
| $MA_{Fri}$ | 0.51 | 0.17 |
| $MA_{Sat}$ | 0.45 | 0.16 |
| $MA_{Sun}$ | 0.81 | -0.03 |
| $EA_{Week}$ | 0.03 | 0.84 |
| $EA_{Fri}$ | -0.03 | 0.84 |
| $EA_{Sat}$ | 0.01 | 0.57 |
| $EA_{Sun}$ | -0.04 | 0.49 |

**S4 Table.** Exploratory bifactor analysis. We perform a bifactor analysis to compute g chronotype described in the main text. The loadings of all the factors, g chronotype, F1* and F2*, factors have been summarised in this table. The general factor g loads directly onto all the calling activities of the individuals, thus computing ascore that is used to identify their chronotypes. F1* and F2* are group factors that load separately onto the morning and the evening activities.

|  | g chronotype | F1* | F2* |
| --- | --- | --- | --- |
| $MA_{Week}$ | 0.49 | 0.74 | |
| $MA_{Fri}$ | 0.37 | 0.42 | |
| $MA_{Sat}$ | 0.34 | 0.38 | |
| $MA_{Sun}$ | 0.43 | 0.68 | |
| $EA_{Week}$ | 0.49 | | 0.70 |
| $EA_{Fri}$ | 0.45 | | 0.70 |
| $EA_{Sat}$ | 0.33 | | 0.48 |
| $EA_{Sun}$ | 0.25 | | 0.40 |

**S5 Table.**
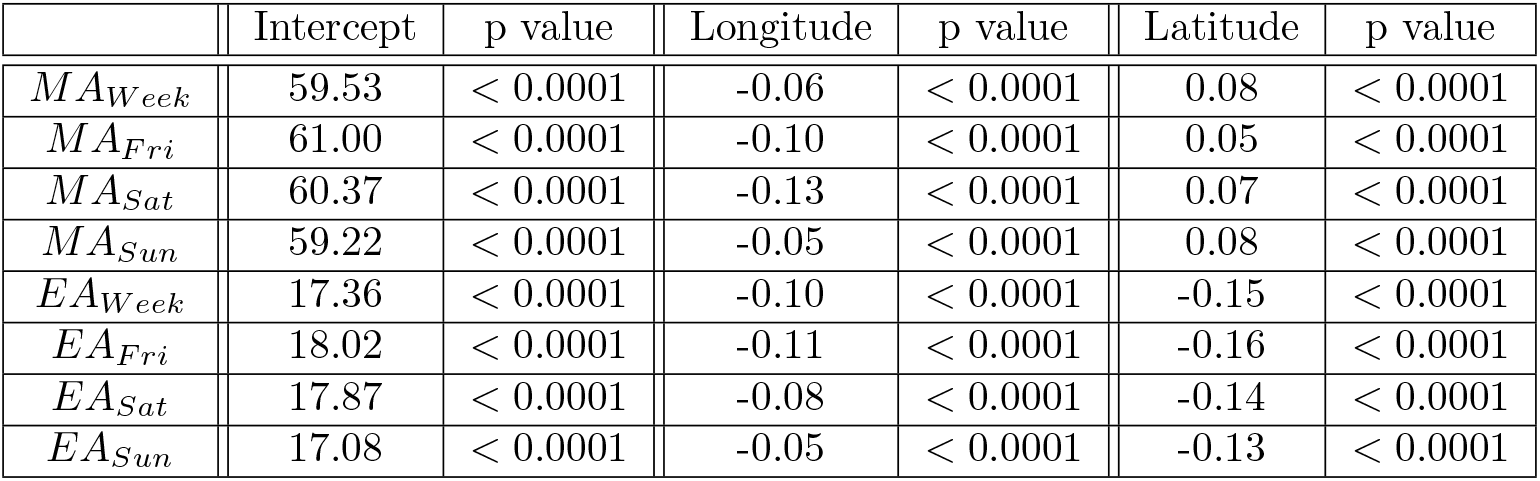
Regression of the calling activities. We again carry out a regression on the morning ({MA_*d*_}) and evening activities ({EA_*d*_}) of the individuals for all d days of the week to remove the geographical effect using longitude and latitude as independent variables. In this table we have summarised the results obtained from computing the regression along with the p values.

|  | Intercept | p value | Longitude | p value | Latitude | p value |
| --- | --- | --- | --- | --- | --- | --- |
| $MA_{Week}$ | 59.53 | < 0.0001 | -0.06 | < 0.0001 | 0.08 | < 0.0001 |
| $MA_{Fri}$ | 61.00 | < 0.0001 | -0.10 | < 0.0001 | 0.05 | < 0.0001 |
| $MA_{Sat}$ | 60.37 | < 0.0001 | -0.13 | < 0.0001 | 0.07 | < 0.0001 |
| $MA_{Sun}$ | 59.22 | < 0.0001 | -0.05 | < 0.0001 | 0.08 | < 0.0001 |
| $EA_{Week}$ | 17.36 | < 0.0001 | -0.10 | < 0.0001 | -0.15 | < 0.0001 |
| $EA_{Fri}$ | 18.02 | < 0.0001 | -0.11 | < 0.0001 | -0.16 | < 0.0001 |
| $EA_{Sat}$ | 17.87 | < 0.0001 | -0.08 | < 0.0001 | -0.14 | < 0.0001 |
| $EA_{Sun}$ | 17.08 | < 0.0001 | -0.05 | < 0.0001 | -0.13 | < 0.0001 |

**S6 Fig.**
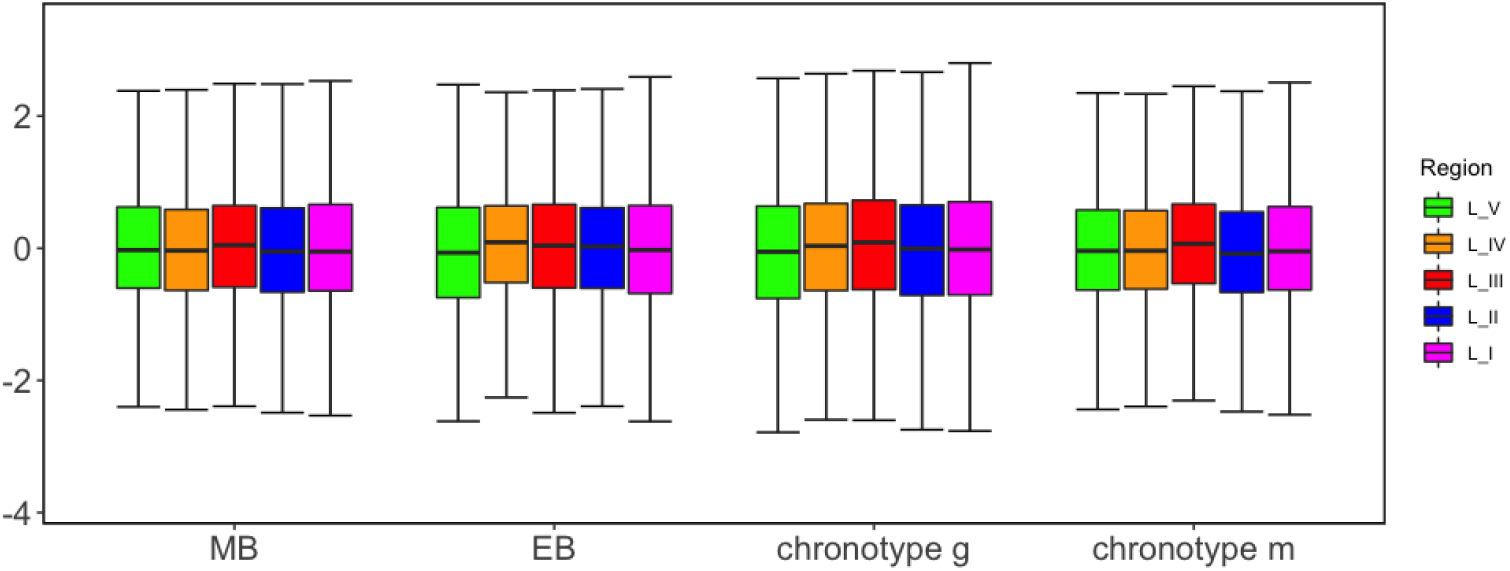
Factor scores of the residuals computed after regression. After carrying out the regression on MAs and EAs we perform an exploratory analysis and bifactor analysis on the residuals obtained. We show the variation of the factor scores of the morning and evening behaviour, the general g chronotype and m chronotype (see S7 Fig) for the 5 regions (longitudinal bands). This plot is computed after an EFA and EBA was done on the residuals to show that we have removed the effect of the East-West progression in the data. Figure shows a summary of the factor scores in the form of a box-plot that includes all the values within the range of the 25^*th*^ and 75^*th*^ percentile and the end of the whiskers represent the maximum and the minimum scores excluding outliers. The horizontal lines inside the boxes represent the median of the scores in each region.

**S7 Fig.**
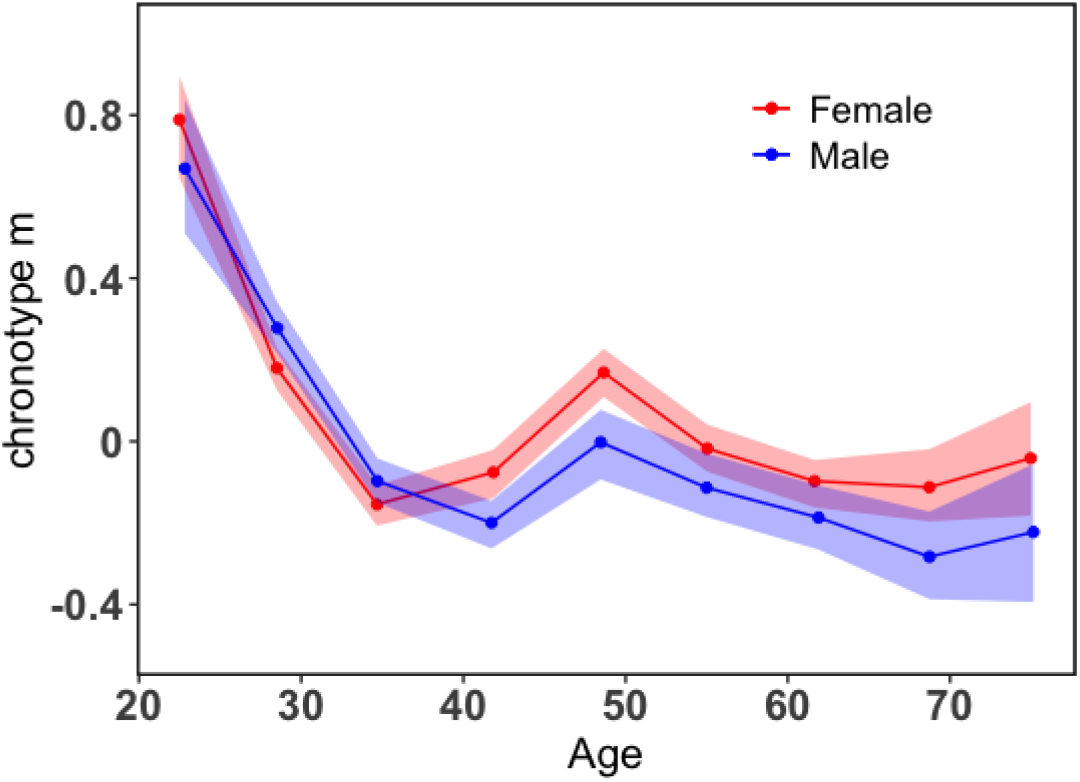
The m chronotype and its variation with age and gender. Finally, we have considered all the 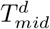 values for all d days of the week. This model has a unidimensionality score is 0.97 which implies that there is only one underlying factor. An EFA carried out on this model also shows that there is one latent factor that can be used as indicator of the chronotype (m chronotype) of an individual as shown in this Figure. The two chronotypes: g and m have a strong correlation (0.78) and both can used to determine the morningness or eveningness of a user. The colour blue has been used males and red for females.

## References

1. Jones, S.E., Lane, J.M., Wood, A.R. et al. Genome-wide association analyses of chronotype in 697,828 individuals provides insights into circadian rhythms. Nat. Commun.. 10, 343 (2019)

2. Merikanto I., Lahti T., Kronholm E., Peltonen M., Laatikainen T., Vartiainen E., Salomaa V. and Partonen T. Evening types are prone to depression. Chronobiology International. 30:5, 719–725 (2013)

3. Facer-Childs E. R., Boiling S. and Balanos G. M. The effects of time of day and chronotype on cognitive and physical performance in healthy volunteers. Sports Medicine - Open. 4:47, (2018)

4. Roenneberg T., Wirz-Justice A. and Merrow M. Life between Clocks: Daily Temporal Patterns of Human Chronotypes. Journal of biological rhythms. 18:1, 80–90 (2003)

5. Roennenberg T. Having Trouble Typing? What on Earth Is Chronotype?. Journal of bilogicat rhythms. 30:6 487–491 (2015)

6. Shahid A., Wilkinson K., Marcu S., Shapiro C.M. (2011) Munich Chronotype Questionnaire (MCTQ). In: Shahid A., Wilkinson K., Marcu S., Shapiro C. (eds) STOP, THAT and One Hundred Other Sleep Scales. Springer, New York, NY.

7. Jung Hie Lee, In Soo Kim, Seong Jae Kim, Wei Wang, Jeanne F Duffy. Change in Individual Chronotype Over a Lifetime: A Retrospective Study. Sleep Med Res. 2, 48–53 (2011)

8. Roenneberg T., Kuehnle T., Pramstaller P. P., Ricken J., Havel M., Guth A., Merrow M. A marker for the end of adolescence. Current Biology. 17:24, R1038 (2004)

9. Adan, A. and Natale, V. Gender differences in morningness-eveningness preference. Chronobiology International. 19, 709–720 (2002)

10. Hotne J. A. and (Östberg O. A self-assessment questionnaire to determine morningness-eveningness in human circadian rhythms. Int. J. Chronobiol. 4(2) 97–110 (1976)

11. Blondel V., de Cordes N., Decuyper A., Deville P., Raguenez J. and Smoreda Z. Mobile Phone Data for Development: Analysis of Mobile Phone Datasets for the Development of Ivory Coast. In S’elected Contributions to the DļD Challenge Sponsored by Orange, MIT, Cambridge, MA, 80–90 (May 1-3, 2013)

12. Eagle N. and Pentland A. Reality mining: sensing complex social systems. Personal and Ubiquitous Computing. 10, 255–268 (2006)

13. Pentland A. Social Physics: How Social Networks Can Make Us Smarter. Penguin, New york. (2015)

14. Onnela J. P., Saramäki J., Hyvönen J., Szabó G., Lazer D., Kaski K., Kertész J. and Barabási A. L. Structure and tie strengths in mobile communication networks. PNAS. 104(18), 7332–7336 (2007)

15. Bhattacharya K. and Kaski K. Social physics: uncovering human behaviour from communication. Advances in Physics: X. 4:1, 1527723 (2018)

16. Miritello G., Moro E., Lara R., Martínez-López R., Belchamber J., Roberts S. G. B. and Dunbar R. I. M. Time as a limited resource: Communication strategy in mobile phone networks. S’ocial Networks. 35:1, 89–95 (2013)

17. Fudolig M. I. D., Bhattacharya K., Monsivais D., Jo H.-H. and Kaski K. Link-centric analysis of variation by demographics in mobile phone communication patterns. PLOS ONE. 15(1) (2020)

18. Fudolig M. I. D., Monsivais D., Bhattacharya K., Jo H.-H. and Kaski K. Different patterns of social closeness observed in mobile phone communication. Journal of Computational Social Science. 3:1 17 (2020)

19. Bhattacharya K., Berg V., Ghosh A., Monsivais D., Kertész J., Kaski K. and Rotkirch A. Different patterns of social closeness observed in mobile phone communication. Journal of Computational Social Science. 3:1 17 (2020)

20. Fudolig M. I. D., Monsivais D., Bhattacharya K., Jo H. H. and Kaski K. Internal migration and mobile communication patterns among pairs with strong ties. arXiv

21. Ghosh A., Berg V., Bhattacharya K., Monsivais D., Kertész J., Kaski K. and Rotkirch A. Migration patterns across the life course of families: Gender differences and proximity with parents and siblings in Finland. https://arxiv.org/pdf/1708.02432.pdf.

22. Grantz K. H., Meredith H. R., Cummings D. A. T., Metcalf C. J. E., Grenfell B. T., Giles J. R., Mehta S., Solomon S., Labrique A., Kishore N., Buckee C. O. and Wesolowski A. The use of mobile phone data to inform analysis of COVID-19 pandemic epidemiology. Nature Communications. 11, 4961 (2020)

23. Monsivais, D., Bhattacharya, K., Ghosh, A., Dunbar, R. and Kaski, K. Seasonal and geographical impact on human resting periods. Scientific Reports. 7, 10717 (2017)

24. Aledavood, T., Lehmann, S. and Saramäki, J. Social network differences of chronotypes identified from mobile phone data. EPJ Data Science. 7, 46 (2018)

25. Aledavood, T., Kivimäki I., Lehmann S. and Saramäki, J. A Non-negative Matrix Factorization Based Method for Quantifying Rhythms of Activity and Sleep and Chronotypes Using Mobile Phone Data. https://arxiv.org/pdf/2009.09914.pdf

26. Roenneberg T., Kuehnle T., Juda M., Kantermann T., Allebrandt K., Gordijn M. and Merrow M. Epidemiology of the human circadian clock. Sleep Medicine Reviews. 11(6) 429–438 (2007)

27. Monsivais, D., Ghosh, A., Bhattacharya, K., Dunbar, R. and Kaski, K. Tracking urban human activity from mobile phone calling patterns. Plos Computational Biology. 13 (2017)

28. There were nine billion calls made in total and one billion text messages sent by all the users in the dataset. So only one-tenth of the total number of the calling activity considered comprise only of text messages. Since the fraction is quite small we do not expect there to be any significant bias in the data for people who prefer texting more than calling.

29. Roenneberg, T., Kumar, C. and Merrow, M. The human circadian clock entrains to sun time. Current Biology. 17, R44–R45 (2007)

30. Reynolds, D. Gaussian Mixture Models.. Encyclopedia Of Biometrics. 741 (2009)

31. Aubourg, T., Demongeot, J., Renard, F., Provost, H. and Vuillerme, N. How to Measure Circadian Rhythms of Activity and Their Disruptions in Humans Using Passive and Unobtrusive Capture of Phone Call Activity.. Studies In Health Technology And Informatics. 264 pp. 1631–1632 (201)

32. Fabrigar L. R., Wegener D. T. Exploratory factor analysis. Oxford University Press, New York. (2012)

33. Costello A. B. and Osborne J. Best practices in exploratory factor analysis: four recommendations for getting the most from your analysis. Practical Assessment, Research, and Evaluation. 10:7 77–85 (2005)

34. Glen S. “Kaiser-Meyer-Olkin (KMO) Test for Sampling Adequacy” From StatisticsHowTo.com: Elementary Statistics for the rest of us! https://www.statisticshowto.com/kaiser-meyer-olkin/

35. Bartlett, M. S. Tests of significance in factor analysis. British Journal of Psychology. 3 77–85 (1950)

36. Revelle W. Package ‘psych’. (2015). https://cran.r-project.org/web/packages/psych/psych.pdf

37. Knekta, E., Runyon, C. and Eddy, S. One size doesn’t fit all: Using factor analysis to gather validity evidence when using surveys in your research. CBE-Life Sciences Education. 18:rm1, 1–17 (2019)

38. Jennrich, R. I. and Bentler, P. M. Exploratory Bi-factor Analysis. Psychometrika. 76(4), 537–549 (2011)

39. Reise, S., Moore, T. and Haviland, M. Bifactor Models and Rotations: Exploring the Extent to which Multidimensional Data Yield Univocal Scale Scores. Journal of Personality Assessment. 92(6), 544–559 (2010)

40. Personality project, http://www.personality-project.org/r/book/chapter7

41. Schmid J., Leiman J. M. The development of hierarchical factor solutions. Psychometrika. 22), 53–61 (1957)

42. Foster R.G. and Roenneberg T. Human Responses to the Geophysical Daily, Annual and Lunar Cycles. Current Biology. 18:17 R784–R794 (2008)

43. Fischer D., Lombardi D.A., Marucci-Wellman H. and Roennenberg T. Chronotypes in the US - Influence of age and sex. PLoS One. 12(6) e0178782 (2017)

44. Ghosh A., Monsivais D., Bhattacharya K., Dunbar R. I. M. and Kaski K. Quantifying gender preferences in human social interactions using a large cellphone dataset. EPJ Data Sci.. 8:9, 89–95 (2019)

45. Duffy J.F., Rimmer D.W. and Czeisler C.A. Association of intrinsic circadian period with morningness–eveningness, usual wake time, and circadian phase. Behav. Neurosci.. 115(4) 895–9 (2001)

46. Henson J., Rowlands A.V., Baldry E., Brady E.M., Davies M.J., Edwardson C.L., Yates T., Hall A.P.; CODEC Investigators. Physical behaviors and chronotype in people with type 2 diabetes. BMJ Open Diabetes Res. Care. 8(1) e001375 (2020).

47. Juda M., Vetter C. and Roenneberg T. Chronotype Modulates Sleep Duration, Sleep Quality, and Social Jet Lag in Shift-Workers. Journal of Biological rhythms. 28:2, 141–151 (2013)

48. Panjeh S., Pompeia S., Archer S. N., Pedrazzoli M., von Schantz M. and Cogo-Moreira H. What are we measuring with the morningness–eveningness questionnaire? Exploratory factor analysis across four samples from two countries. Chronobiology International. (2020)

49. Onnela J. P. and Rauch S. L. Harnessing smartphone-based digital phenotyping to enhance behavioral and mental health. Neuropsychopharmacology. 41(7), 1691 (2016)

50. Huguet A., Rao S., McGrath P. J., Wozney L., Wheaton M., Conrod J. and Rozario S. A systematic review of cognitive behavioral therapy and behavioral activation apps for depression. PLoS ONE. 11(5), e0154248 (2016)

